# SARS-CoV-2 utilizes a multipronged strategy to suppress host protein synthesis

**DOI:** 10.1101/2020.11.25.398578

**Authors:** Yaara Finkel, Avi Gluck, Roni Winkler, Aharon Nachshon, Orel Mizrahi, Yoav Lubelsky, Binyamin Zuckerman, Boris Slobodin, Yfat Yahalom-Ronen, Hadas Tamir, Igor Ulitsky, Tomer Israely, Nir Paran, Michal Schwartz, Noam Stern-Ginossar

## Abstract

Severe acute respiratory syndrome coronavirus 2 (SARS-CoV-2) is the cause of the ongoing coronavirus disease 19 (COVID-19) pandemic. Despite the urgent need, we still do not fully understand the molecular basis of SARS-CoV-2 pathogenesis and its ability to antagonize innate immune responses. Here, we use RNA-sequencing and ribosome profiling along SARS-CoV-2 infection and comprehensively define the mechanisms that are utilized by SARS-CoV-2 to shutoff cellular protein synthesis. We show SARS-CoV-2 infection leads to a global reduction in translation but that viral transcripts are not preferentially translated. Instead, we reveal that infection leads to accelerated degradation of cytosolic cellular mRNAs which facilitates viral takeover of the mRNA pool in infected cells. Moreover, we show that the translation of transcripts whose expression is induced in response to infection, including innate immune genes, is impaired, implying infection prevents newly transcribed cellular mRNAs from accessing the ribosomes. Overall, our results uncover the multipronged strategy employed by SARS-CoV-2 to commandeer the translation machinery and to suppress host defenses.

## Introduction

Severe acute respiratory syndrome coronavirus 2 (SARS-CoV-2) is the cause of the ongoing coronavirus disease 19 (COVID-19) pandemic ^1,2^. SARS-CoV-2 is an enveloped virus consisting of a positive-sense, single-stranded RNA genome of ~30 kb. From the positive strand genomic RNA, two overlapping open reading frames (ORFs) are translated, ORF1a (pp1a) and ORF1b (pp1ab). The translation of ORF1b is mediated by a −1 frameshift that allows translation to continue beyond the stop codon of ORF1a. These generate continuous polypeptides which are cleaved into a total of 16 nonstructural proteins (NSP 1-16, Snijder et al., 2016; Sola et al., 2015; Stadler et al., 2003; V’kovski et al., 2020). Among them are the subunits that constitute the RNA-dependent RNA polymerase (RdRP), which in turn transcribes subgenomic RNAs that contain a common 5’ leader fused to different segments from the 3′ end of the viral genome ^3–5^. The different subgenomic RNAs encode 4 conserved structural proteins- spike protein (S), envelope protein (E), membrane protein (M), nucleocapsid protein (N)- and several accessory proteins and non-canonical protein products ^7–9^.

Translation of viral proteins relies solely on the cellular translation machinery; therefore, viruses must commandeer this machinery to translate their own mRNAs. In addition, viruses need to counteract the antiviral response cells mount upon infection ^10^. The first line of defense applied by infected cells engages the interferon (IFN) pathway, which amplifies signals resulting from detection of intracellular viral components to induce a systemic anti-viral response. Specifically, cells contain various sensors that detect the presence of viral RNAs and promote nuclear translocation of transcription factors, leading to transcription and secretion of type I IFNs ^11^. Binding of IFN to its cognate receptor in autocrine and paracrine manners leads to the propagation of the signal and to the transcription and translation of hundreds of interferon stimulated genes that act to hamper viral replication at various stages of the viral life cycle ^12^. In the case of SARS-CoV-2, interferon response seems to play a critical role in pathogenesis ^13–17^. In addition, the extent to which SARS-CoV-2 suppresses the IFN response is a key characteristic that distinguishes it from other respiratory viruses ^18,19^.

Viruses utilize various strategies to cause shutoff of host mRNA translation ^10^, including hampering mRNA processing steps and export, inducing degradation of mRNAs and inhibiting translation. Coronaviruses (CoVs) are known to cause host shutoff ^10,20^ and several strategies have been proposed for how the beta CoVs may shut-off host protein synthesis and evade immune detection. These include degradation of host mRNA in the nucleus or the cytosol and inhibition of host translation ^21,22^. Nonetheless, the extent to which SARS-CoV-2 uses these or other strategies remains unclear.

NSP1 is the best characterized and most prominent coronavirus host shutoff factor ^23^. Several recent studies showed SARS-CoV-2 NSP1 binds the 40s ribosome and inhibits translation ^24–28^. In addition, other SARS-CoV-2 proteins were shown to interfere with cellular gene expression. For example, one of the SARS-CoV-2 accessory proteins, ORF6, was suggested to disrupt nucleocytoplasmic transport leading to inhibition of gene expression ^29^ and several viral proteins have been demonstrated to antagonize IFN-I production and signaling ^24,30^. Although, different SARS-CoV-2 proteins are implicated in host expression shutoff, a comprehensive picture of the effect of SARS-CoV-2 infection on cellular gene expression is still lacking.

Here we employ RNA-sequencing and ribosome profiling along SARS-CoV-2 infection to explore the mechanisms which the virus utilizes to interfere with host protein synthesis. We reveal SARS-CoV-2 uses a multi-faceted approach to shutoff cellular protein production. SARS-CoV-2 infection induces global translation inhibition but surprisingly the translation of viral transcripts is not preferred over their cellular counterparts. Instead we reveal that infection leads to accelerated cellular mRNAs degradation, likely conferred by NSP1. Viral transcripts are refractory to these effects, an evasion potentiated by their 5’UTRs, enabling viral dominance over the mRNA pool in infected cell. Finally, we show that the translation of transcripts whose expression is induced in response to infection, including innate immune genes, is severely impaired. Overall these findings reveal the key mechanisms SARS-CoV-2 is applying to suppress cellular protein synthesis and host defenses.

## Results

### Simultaneous monitoring of RNA levels and translation during SARS-CoV-2 infection

To gain a detailed view of the changes that occur in viral and host translation over the course of SARS-CoV-2 infection, we infected Calu3 cells with SARS-CoV-2 at multiplicity of infection (MOI) of 3 and harvested infected cells at 3, 5, and 8 hours post infection (hpi) as well as uninfected cells. This high MOI infection results in infection of the majority of the cells and thus a synchronous cell population, allowing for molecular dissection of the events along infection. We designed our experiment to simultaneously monitor both RNA levels and translation (Figure 1A). Deep sequencing of mRNA (RNA-seq) generates a detailed depiction of transcript levels during infection and this was paired with ribosome profiling, which allows accurate measurement of protein synthesis by capturing the overall *in-vivo* distribution of ribosomes on a given message ^31,32^. In order to assess the reproducibility of our experiments we prepared two independent biological replicates for the uninfected and 8hpi time points and both the mRNA and footprint measurements were reproducible (Figure S1A and S1B). Metagene analysis, in which gene profiles are aligned and then averaged, revealed the expected profiles of footprints along mRNAs; ribosome density accumulates from the start codon along the gene body ending at the first in-frame stop codon with pronounced accumulation of ribosomes at the initiation sites and 3 bp periodicity along the gene body (Figure S1C). Using this data, we quantitatively assessed the expression pattern of 8627 cellular transcripts and 12 canonical viral ORFs that are expressed from the genomic and sub-genomic RNAs along SARS-CoV-2 infection.

**Figure 1:**
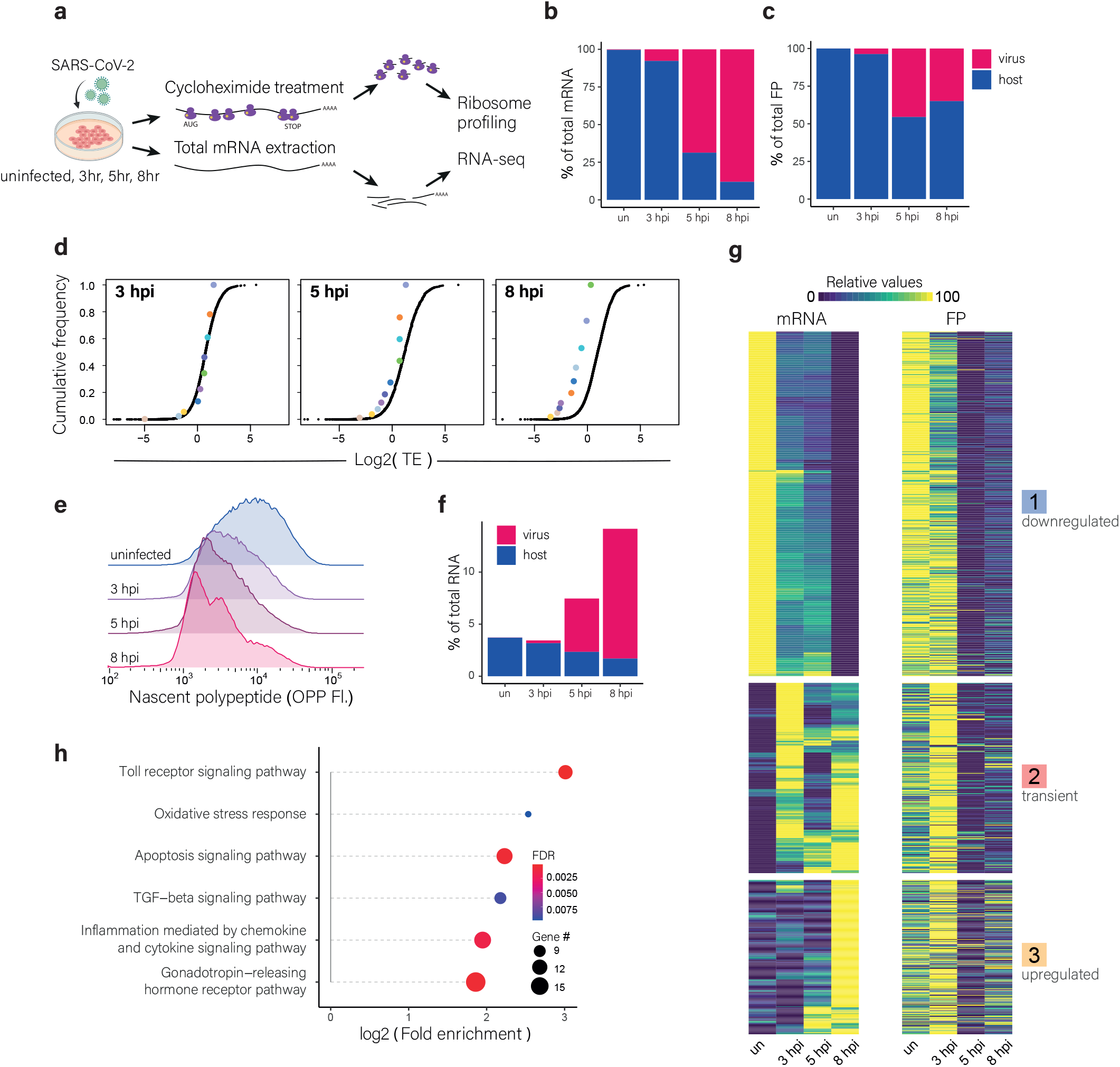
Global reduction of translation and in cellular mRNA levels along SARS-CoV-2 Infection **(A)** Calu3 cells were left uninfected or were infected with SARS-CoV-2 (MOI=3) for 3, 5 or 8 hours and harvested for RNA-seq and for Ribo-seq. **(B and C)** Percentage of reads that aligned to human or viral transcripts from the sum of aligned reads shown for mRNAs **(B)** and footprints **(C)** in uninfected cells and in cells harvested at 3, 5 and 8 hpi. **(D)** Cumulative TE distribution among well-expressed human (black points) and viral (colored points) genes at 3, 5 and 8 hpi. **(E)** Protein synthesis measurement by flow cytometry of Calu3 cells infected with SARS-CoV-2 (MOI = 3) for 3, 5 and 8 hpi or an uninfected control following O-Propargyl Puromycin (OPP) incorporation and fluorescent labelling. **(F)** Percent of reads that aligned to the human or viral transcripts from the sum of total RNA reads in uninfected cells and in cells harvested at, 3, 5 and 8 hpi. **(G)** Heat map presenting relative mRNA and footprints expression of well-expressed human transcripts that showed the most significant changes in their mRNA levels at 8 hpi relative to uninfected, across time points during SARS-CoV-2 infection. Shown are expression levels scaled by gene after partitioning clustering. Three main clusters are marked on the right. **(H)** Summary of pathway enrichment analysis of genes enriched in cluster 3 (upregulated genes). Dot size reflects the number of genes from each pathway included in the tested set, and dot color reflects the false discovery rate (FDR) of the pathway enrichment.

Analysis of the mRNAs and footprints originating from cellular and viral transcripts illustrates SARS-CoV-2’s unprecedented dominance over the mRNA pool. At 8 hpi viral mRNAs comprise almost 90% of the mRNAs in infected cells (Figure 1B). Surprisingly, however, at the same time point, viral mRNAs account for only ~34% of the RNA fragments engaged with ribosomes in the cells (Figure 1C). In order to quantitatively evaluate the ability of SARS-CoV-2 to co-opt the host ribosomes we calculated the relative translation efficiency (TE) of viral and cellular RNA along infection. TE is defined as the ratio of footprints to mRNA reads for a given gene and reflects how well a gene is being translated. We then compared the TE of human genes to that of viral genes at each of the time points along infection. The analysis shows that at 3 hpi viral gene translation efficiencies fall within the general range of cellular gene translation. This indicates that when infection initiates, viral transcripts are translated with efficiencies similar to those of host transcripts. As infection progresses, viral gene translation efficiency relative to cellular genes is significantly reduced (Figure 1D). Since it seems improbable that the virus will promote conditions that are unfavorable to the translation of its mRNA, this relative reduction in translation of viral genes at 5 and 8 hpi may indicate that not all viral RNAs are accessible for translation. Since double-membrane sealed replication compartments are formed to accommodate viral genome replication ^33^, an appealing possibility is that these compartments sequester a sizable fraction of the viral RNA molecules and thus prevent them from being a part of the translated mRNA pool.

By their nature, deep sequencing measurements provide relative values but not absolute quantification of RNA and translation levels. Since SARS-CoV-2 encoded protein, NSP1, was recently shown to interfere with translation by blocking the mRNA entry channel of ribosomes ^25–28^, and since the extent to which SARS-CoV-2 interferes with the overall levels of cellular mRNA was not assessed, we next examined if SARS-CoV-2 infection affects global translation and RNA levels. To quantify absolute translation levels, we measured nascent protein synthesis levels using an analogue of puromycin, O-Propargyl Puromycin (OPP), which incorporates into elongating polypeptide chains ^34^. The nascent polypeptides with incorporated OPP are fluorescently-labeled via a Click reaction and the levels of OPP incorporation into elongating polypeptides is quantified by flow cytometry (Figure S2A and S2B). We infected Calu3 cells with SARS-CoV-2 at an MOI of 3 and measured nascent protein synthesis levels in uninfected cells and at 3, 5 and 8 hpi. We observed a significant reduction in global translation levels already at 3 hpi which was augmented with time, and at 8 hpi translation activity was reduced by 50% (Figure 1E). In parallel we measured the total RNA levels and rRNA levels extracted from uninfected cells and at 3, 5, and 8 hpi. This analysis illustrated there are no major changes in total RNA or in rRNA levels along SARS-CoV-2 infection (Figure S3A and S3B). Since the vast majority of RNA in cells originates from rRNA and this dominance of rRNA may mask changes in mRNA levels we sequenced total RNA, without rRNA depletion, to assess the relative abundance of cellular and viral mRNAs in uninfected cells and at 3, 5, and 8 hpi. This analysis demonstrates that the pool of mRNA molecules relative to rRNA is growing during infection, due to the massive production of viral transcripts but at the same time the relative fraction of cellular mRNA is reduced by approximately 2-fold (Figure1F). This suggests that during infection there is both massive production of viral transcripts and a concomitant substantial reduction in the levels of cellular transcripts.

Next, we quantitatively assessed the expression pattern of cellular genes along SARS-CoV-2 infection. We clustered the mRNA levels of genes that showed the most significant changes along infection using partitioning clustering, allowing grouping of cellular transcripts into three distinct classes based on similarities in temporal expression profiles in the RNA-seq. Overall, we found that changes in ribosome footprints tracked the changes in transcript abundance (Figure 1G), with some exceptions that will be discussed in detail below. This shows that part of the reduction in cellular protein synthesis is driven by the reduction in cellular RNA levels. Interestingly, although the levels of the majority of host transcripts were reduced during SARS-CoV-2 infection, we identified numerous transcripts that were significantly elevated (Figure 1G, cluster 3). We carried out pathway enrichment analysis for each of these three clusters. As expected, the group of upregulated mRNAs (cluster 3) was significantly enriched with genes related to immune response, including Toll receptor Signaling, chemokine and cytokine signaling (Figure 1H and Table S1) and these upregulated genes include IL6 and IL8 which play a significant role in the pathogenesis of SARS-CoV-2 ^35^ and several IFN stimulated genes like, IFIT1, 2 and 3, IRF1, ISG15 and TNF alpha induced proteins. These measurements and analyses reveal that the shutoff in host protein synthesis is driven by several mechanisms including; general reduction in the translation capacity of infected cells and reduction in the levels of most cellular mRNAs.

### Viral gene expression dynamics along infection

We next analyzed viral gene expression dynamics along SARS-CoV-2 infection. Viral ORFs are translated from the 30kb genomic RNA or from a nested series of subgenomic RNAs that contain a common 5′ leader fused to different segments from the 3′ end of the viral genome (Figure 2A and V’kovski et al., 2020). Viral transcripts and translation levels significantly increased from 3 to 5 hpi but the relative abundance of some viral transcripts and their translation rates were reduced from 5 to 8 hpi (Figure 2B and 2C). This relative reduction in translation rates of some viral transcripts prompted us to assess the relative translation efficiency of viral genes along infection. Since, as indicated above, the translation efficiency of viral genes compared to their cellular counterparts is relatively reduced along infection (Figure 1D), here we examined how the translation of viral genes is distributed between different viral transcripts at different times post infection. This focused analysis revealed an interesting connection between changes in translation efficiency of viral ORFs and their relative location on the viral genome. ORFs that are located at the 5’ end of the genome showed relative increase in their translation efficiency along infection whereas ORFs that are encoded towards the 3’ end of the genome showed relative reduction in their translation efficiencies and ORFs located in the middle of genome showed no major changes in their relative translation efficiency (Figure 2D). Since all viral subgenomic RNAs share the same 5’UTR these differences in translation capacity according to the location in the genome point to a potential unappreciated role for the 3’UTR (the 3’ portion of which is shared between all transcripts) or for the length of viral transcripts, which varies greatly between viral transcripts (Figure 2A). To examine if this phenomenon reflects a general beta CoVs feature, we analyzed the expression and translation of Mouse Hepatitis Virus strain A59 (MHV) ORFs along MHV infection using published RNA-seq and ribosome profiling data along MHV infection ^36^. MHV has a similar genomic organization (Figure 2E) and the increase in viral mRNA and translation between 5 and 8 hpi varied between viral ORFs (Figure 2F and 2G). Although this phenomenon was not as strong, also in the case of MHV, ORFs that are located at the 5’ end of the genome showed relative increase or no change in their translation efficiency along infection whereas ORFs that are encoded towards the 3’ end of the genome showed relative reduction in their translation efficiency (Figure 2H). Together these results indicate that the 3’UTR or viral gene length are likely regulating the translation efficacy of viral mRNAs along infection.

**Figure 2:**
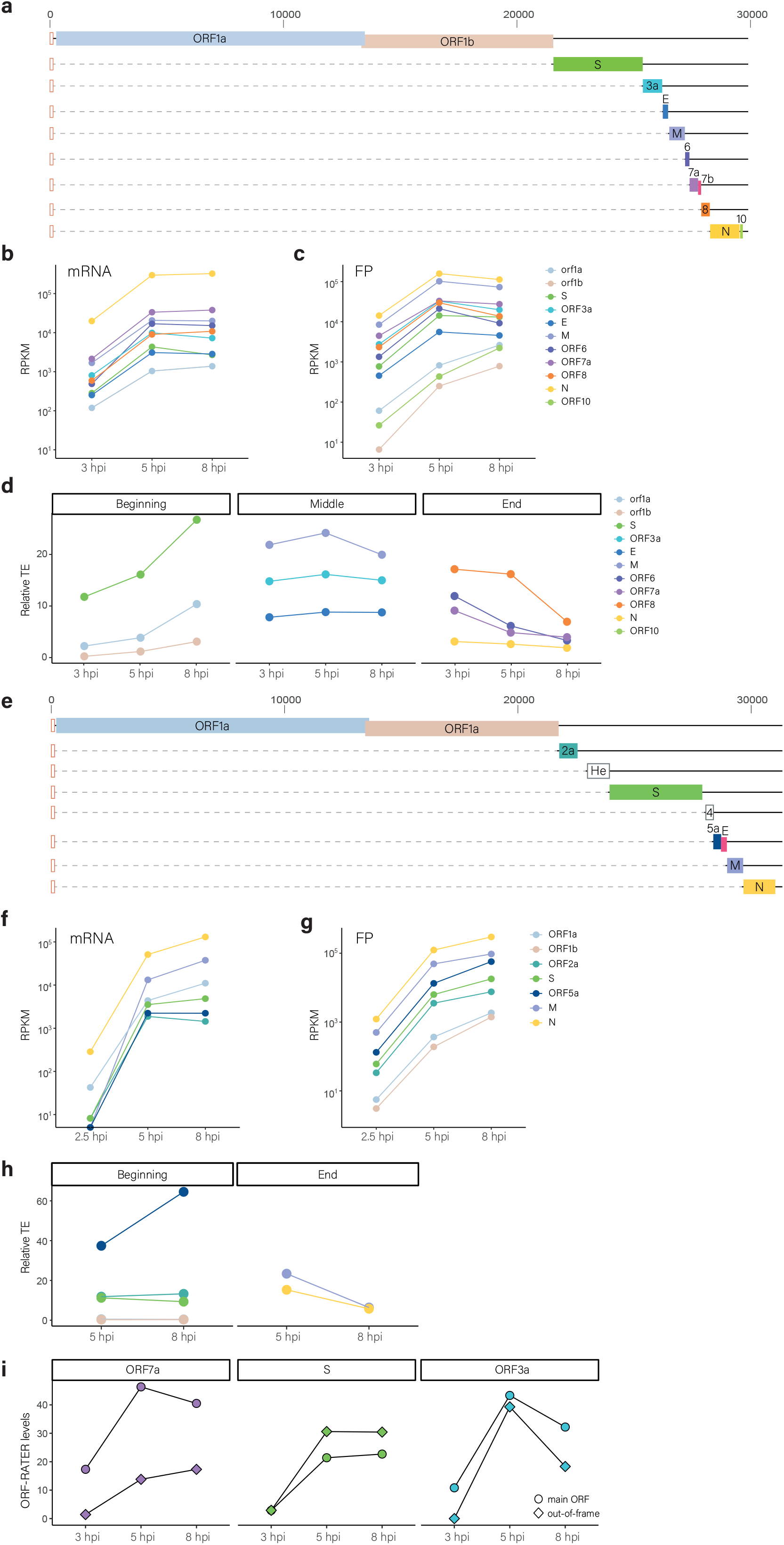
Changes in viral gene expression along SARS-CoV-2 and MHV infection **(A)** Schematic presentation of the SARS-CoV-2 genome organization, the subgenomic mRNAs and the main ORFs. **(B and C)** RNA **(B)** and translation levels **(C)** of each of the SARS-CoV-2 canonical ORFs at 3, 5 and 8 hpi. **(D)** Relative translation efficiency of each canonical viral ORF along infection. Genes are divided to three groups based on their physical location along the genome. **(E)** Schematic presentation of the MHV subgenomic mRNAs, and the main ORFs. ORFs that are not expressed in the strain that was used for ribosome profiling (MHV-A59) are marked by empty rectangles. (**F and G)** RNA **(F)** and translation levels **(G)** of each of the MHV ORFs at 2.5, 5 and 8 hpi. Expression was calculated from ^36^. **(H)** Relative translation efficiency of each canonical viral ORF along infection. Genes are divided to two groups based on their location on the genome. **(I)** Relative translation efficiency of the main ORFs (ORF7a, S and ORF3a, labeled by circle) and the out of frame ORFs (ORF7b, S.iORF and ORF3c, labeled by diamond) of the S, ORF3a and ORF7a subgenomic transcripts, respectively, along SARS-CoV-2 infection. Translation levels were calculated from ribosome densities using ORF-RATER ^49^.

Most SARS-CoV-2 ORFs are 3′-proximal and translated from dedicated subgenomic mRNA (Figure 2A). However several subgenomic RNAs encode for additional out-of-frame ORFs, likely via a leaky scanning mechanism ^9,37–39^. These include ORFs 7b and ORF9b, which are translated from the ORF7a and N subgenomic RNAs, and two ORFs we recently identified by ribosome profiling ^9^, that are supported by additional evidence ^37,39^, ORF3c and ORFS.iORF that are translated from ORF3a and ORF-S subgenomic RNAs. Since scanning efficiency can be regulated by stress conditions ^40^ we examined if the ratio between the translation of a 3′-proximal ORF and its corresponding out-of-frame ORF (encoded by the same subgenomic RNA) changes during infection. Since ORF9b expression was low in our measurements it was excluded from this analysis. The translation of ORF7b, ORF3c and ORFS.iORF correlated with the expression of the 3′-proximal main ORF, indicating there are no major changes in the efficiency of ribosome scanning of viral transcripts along infection (Figure 2I).

### Cellular mRNAs are degraded during SARS-CoV-2 infection

Our results indicate that the levels of the majority of cellular RNA are reduced during SARS-CoV-2 infection and this reduction contributes to the shutoff of cellular protein synthesis. Reduction in cellular RNA levels could be due to interference with RNA production or accelerated RNA degradation. To explore the molecular mechanism, we analyzed if the reduction in cellular transcripts is associated with their subcellular localization. We used measurements of the subcellular localization of transcripts by cytoplasmic and nuclear fractionation^41^ to assess the importance of subcellular localization. The levels of transcripts that mostly localize to the cytoplasm were more reduced in infected cells compared to transcripts that are mostly nuclear (Figure 3A) and there was a clear correlation between subcellular localization and the extent of reduction in transcript levels following SARS-CoV-2 infection (Figure S4A). Furthermore, compared to transcripts encoded in the nuclear genome, mitochondrial encoded transcripts were refractory to the effects of SARS-CoV-2 infection (Figure 3B). The specific sensitivity of cytosolic transcripts implies these transcripts may be specifically targeted during SARS-CoV-2 infection. In CoVs, the most prominent and well characterized cellular shutoff protein is NSP1 ^23^. So far, studies on SARS-CoV-2 NSP1 demonstrated it restricts translation by directly binding to the ribosome 40S subunit ^25–28^, thereby globally inhibiting translation initiation. For SARS-CoV, on top of this translation effect, NSP1 interactions with the 40S was also shown to induce cleavage of translated cellular mRNAs, thereby accelerating their turnover ^21^. To examine if the reduction in cellular transcripts in SARS-CoV-2 infected cells is coupled to their translation we compiled a list of 14 long non-coding RNAs (lncRNAs) that localize to the cytoplasm and are well expressed in our data but, as expected, poorly translated (Figure S4B). Relatively to cellular mRNAs, cytoplasmic lncRNAs were less affected by SARS-CoV-2 infection (Figure 3C), indicating accelerated turnover of cellular transcripts in infected cells may be related to their translation. Recently, ribosome profiling and RNA-seq were conducted on cells transfected with NSP1 ^42^. Analysis of the RNA expression from this data revealed that ectopic NSP1 expression leads to weaker but similar signatures to the ones we identified in infected cells; stronger reduction of cytosolic transcripts compared to nuclear transcripts, stronger sensitivity of nuclear encoded transcripts, and stronger reduction of translated mRNA compared to cytosolic lncRNAs (Figure S5A-C).

**Figure 3:**
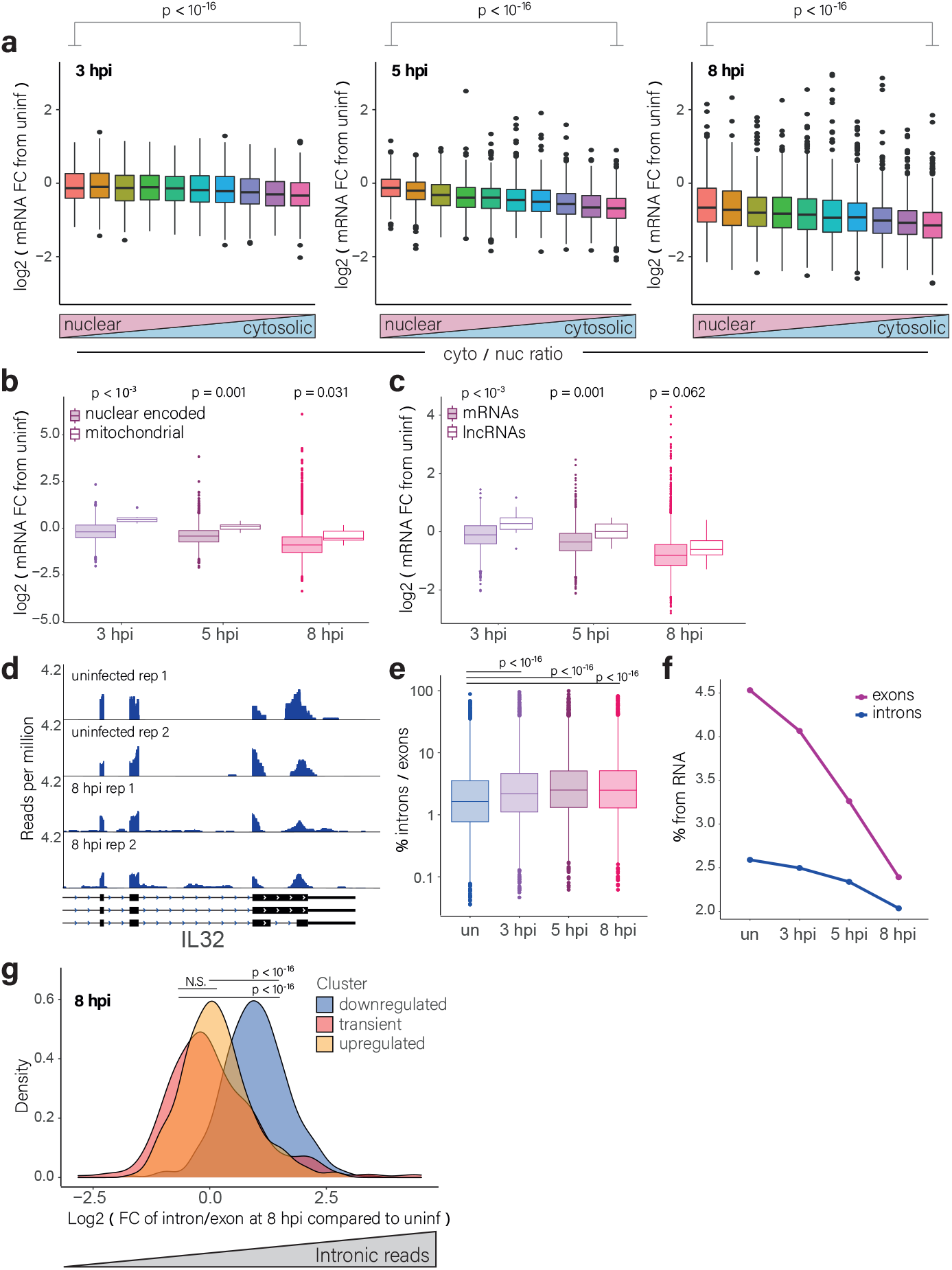
Cytosolic cellular RNAs are degraded during SARS-CoV-2 infection **(A)** RNA level fold change of cellular RNAs at different time points after infection relative to uninfected cells. RNAs were grouped to ten bins based on their cytosol to nucleus localization ratio ^41^. **(B)** The change in RNA levels of nuclear encoded or mitochondrial encoded RNAs at different time points after infection relative to uninfected cells. **(C)** The change in RNA expression of cytosolic lncRNAs and protein coding mRNAs at different time points after infection relative to uninfected cells. **(D)** RNA reads on exons and introns of the end of IL-32 gene from uninfected cells and at 8 hpi. **(E)** Box plots presenting the ratio of intronic to exonic reads for each gene in uninfected cells and at the different time points along SARS-CoV-2 infection. **(F)** The % of reads that align to exonic or intronic regions relative to rRNA abundance along SARS-CoV-2 infection. **(H)** Histograms showing the change in the ratio of intronic to exonic reads of cellular genes at 8hpi relative to uninfected cells. Genes are divided according to the three clusters shown in figure 1G (representing different expression pattern along infection).

We further noticed SARS-CoV-2 infection leads to increased levels of intronic reads in many cellular transcripts (Figures 3D and 3E) indicating SARS-CoV-2 may interfere with cellular mRNA splicing, as was recently suggested ^24^. However, massive degradation of mature cytosolic mRNAs may also generate a relative increase in intronic reads. We therefore analyzed the ratio of intronic and exonic reads relative to rRNA. Whereas relative to rRNA levels, exonic reads showed drastic reduction along SARS-CoV-2 infection, the intronic reads levels showed a more subtle change (Figure 3F). Furthermore, the increase in the ratio of intronic to exonic reads was greater in genes whose expression was reduced along infection compared to genes whose expression was induced (Figure 3G and Figure S6A), illustrating the relative increase in intronic reads is mostly independent of newly transcribed RNAs. Finally, we also detected more intronic reads in cells that exogenously expressed NSP1^42^ (Figure S6B). Together these results indicate that the increase in intronic reads compared to exonic reads during SARS-CoV-2 infection is largely driven by accelerated degradation of mature cellular transcripts that leads to relative reduction in exonic reads. It is likely that SARS-CoV-2 also directly regulates splicing efficiency, as was recently proposed ^24^, but this effect seems more subtle. Together these findings indicate SARS-CoV-2 infection leads to accelerated degradation of cytosolic cellular mRNAs and that this degradation significantly affects the production of cellular proteins.

### SARS-CoV-2 5’UTR protects viral RNA from NSP1 induced RNA degradation

An important aspect of host shutoff during infection is the ability of the virus to hamper the translation of cellular transcripts while recruiting the ribosome to its own transcripts. Although it has been suggested that SARS-CoV-2 mRNAs are refractory to the translation inhibition induced by NSP1 ^24,43^ our measurements indicate that SARS-CoV-2 transcripts are not preferentially translated in infected cells (Figure 1D and ^9^), that RNA degradation plays a prominent role in remodeling the mRNA pool in infected cells and that SARS-CoV-2 dominates the mRNA pool. All SARS-CoV-2-encoded subgenomic RNAs contain a common 5′ leader sequence that is added during negative-strand synthesis ^44^. We therefore explored whether the 5’UTR sequence protects viral mRNAs from NSP1 induced degradation. We fused the viral 5’UTR sequence or a short control 5’UTR to the 5′ end of a GFP reporter (Figure 4A) and transfected these constructs together with expression vectors encoding NSP1 or NSP2 (that was used as a control) into 293T cells. We found that NSP1 expression suppresses the production of the control-GFP but not of the 5’UTR-containing GFP (Figure 4B and 4C). We extracted RNA from these cells and observed that the NSP1 induced reduction in control-GFP expression was associated with 15-fold reduction in the RNA GFP levels whereas the levels of the GFP RNA with the SARS-CoV-2 5’UTR were not reduced, and were even slightly increased, by NSP1 expression (Figure 4D). The plasmid we used also contains an mCherry reporter expressed from an independent promoter. Reassuringly NSP1 also induces a reduction in both mCherry protein (Figure S7A and S7B) and RNA levels (Figure S7C) when compared to NSP2. These results indicate that the 5′ UTR of viral RNAs provides them protection from NSP1 induced degradation and that this protection contributes to the ability of the virus to dominate the mRNA pool in infected cells.

**Figure 4:**
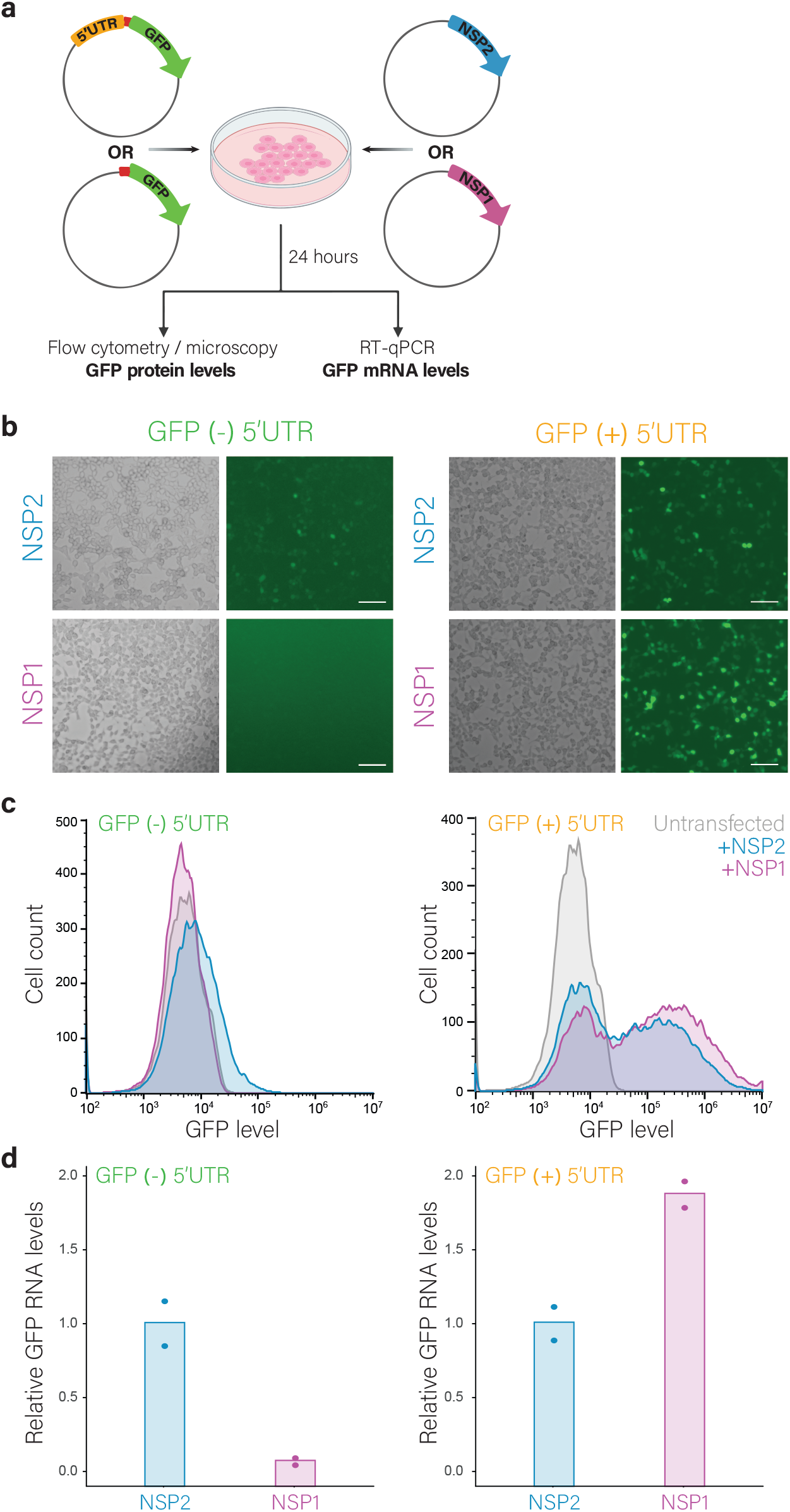
The viral 5′ UTR protects mRNA from NSP1-mediated degradation **(A)** 293T cells were transfected with expression vectors encoding either NSP1 or NSP2 (as a control) and with a GFP reporter (GFP (−) 5’UTR) or a GFP reporter that includes the viral 5’UTR (GFP (+) 5’UTR). **(B)** Microscopy images of cells co-transfected with NSP2 (top) or NSP1 (bottom) together with either GFP (−) 5’UTR or GFP (+) 5’UTR. Scale bars are 100μm. **(C)** Flow cytometry analysis of cells co-transfected with NSP1 or NSP2 together with either GFP (−) 5’UTR or GFP (+) 5’UTR. **(D)** Relative GFP RNA levels from GFP (−) 5’UTR or GFP (+) 5’UTR in cells expressing NSP1 or NSP2 as measured by quantitative RT-PCR. Data points show measurement of biological replicates. Shown is one representative experiment out of two performed.

### The translation efficiency of transcriptionally induced genes is impaired during infection

Our results so far exemplify how SARS-CoV-2 remodels the transcript pool in infected cells. To quantitatively evaluate the role of translational control along SARS-CoV-2 infection, we calculated relative translation efficiency (TE, ratio of footprints to mRNAs for a given gene) along infection. We then centered on genes that showed the strongest change in their relative TE along infection. We clustered these genes into four clusters, based on similarity in their temporal TE profiles, which largely reflect genes whose relative TE is reduced along infection and genes whose relative TE is increased. The mRNA and footprint temporal profiles of these genes reveal a clear signature; the genes whose relative TE along infection was reduced were genes whose mRNA increased during infection but did not show a corresponding increase in footprints (Figure 5A and Figure S8). These genes were enriched in immune response genes (FDR < 10^−4^) like IRF1, IL-6 and CXCL3. Comparing changes in mRNA and TE levels of cellular genes along infection demonstrates that generally transcripts which are transcriptionally induced following infection show a reduction in their relative TE and vice versa (Figure 5B). This inability of RNA that is elevated in response to infection to reach the ribosomes, may explain why infected cells fail to launch a robust IFN response ^13,14^. These data demonstrate that the overall capacity of infected cells to generate new proteins is severely impaired.

**Figure 5:**
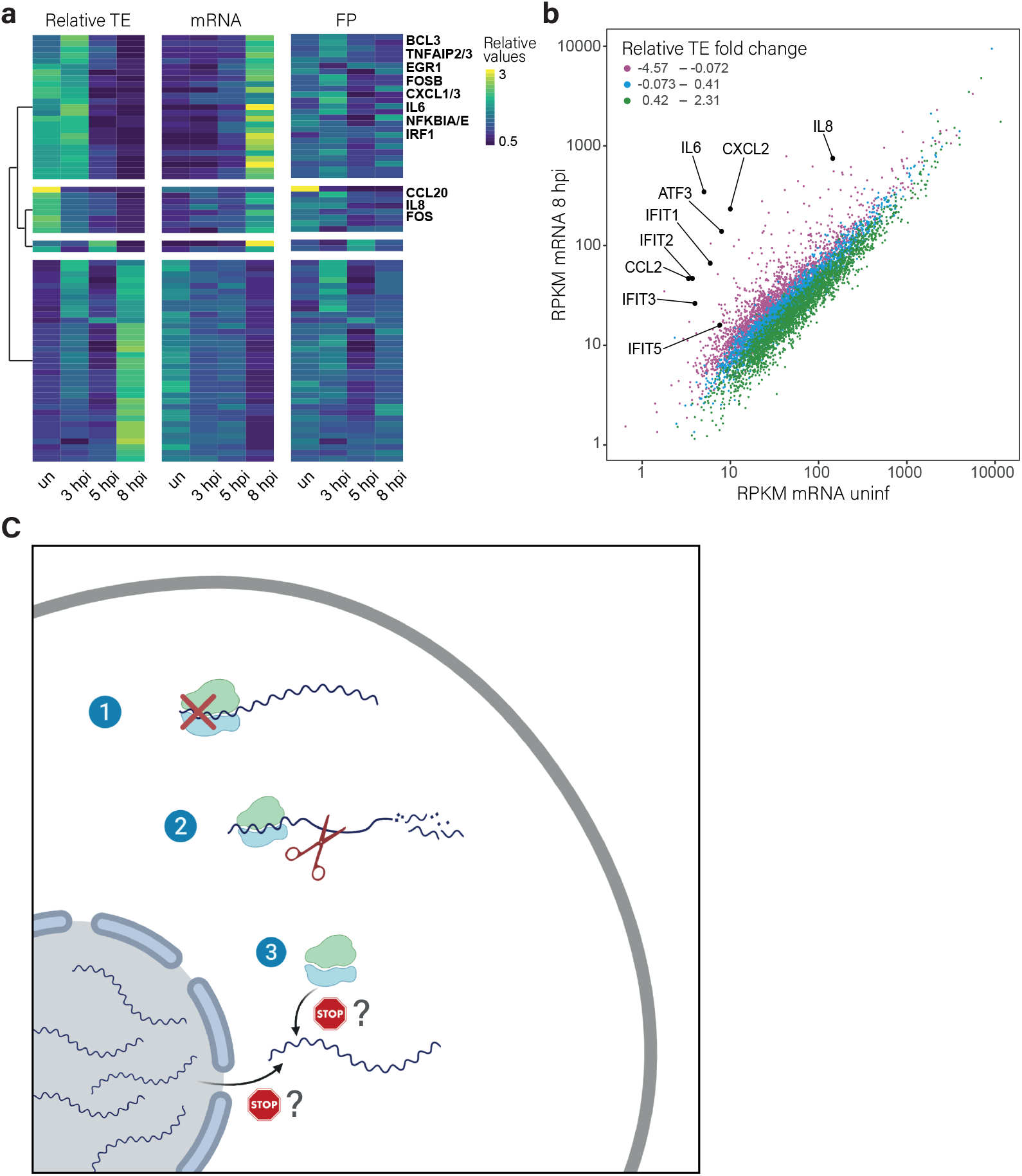
The translation of induced transcripts is impaired during infection **(A)** Heat map presenting relative TE, mRNA and footprints (FP) of human genes that showed the most significant changes in their relative TE along SARS-CoV-2 infection. Shown are relative expression ratios after partitioning clustering based on changes in relative TE values. **(B)** Scatter plot presenting cellular transcript levels in uninfected cells compared to 8hpi. Genes are colored based on the relative change in their TE between uninfected and 8hpi. Central cytokines and IFN stimulated genes are labeled. **(C)** A model of how SARS-CoV-2 suppresses host gene expression through multi-pronged approach: 1. Global translation reduction; 2. Degradation of cytosolic cellular mRNAs; 3. Specific translation inhibition of newly synthesized cellular mRNAs.

## Discussion

Many viruses developed varied and sophisticated mechanisms to repress host mRNA translation in order to block the cellular innate immune response and concomitantly allow the translation of viral mRNAs. Specifically, CoVs have evolved specialized mechanisms to hijack the host gene expression machineries, including inhibition of host protein synthesis and induction of endonucleolytic cleavage of host mRNAs ^23,45^. In the case of SARS-CoV-2 several viral ORFs have been suggested to interfere with viral gene expression ^24–28,30^, but what occurs during viral infection remained an open question.

Using unbiased measurements of translation and RNA expression along SARS-CoV-2 infection, we identified three major courses by which SARS-CoV-2 interferes with cellular gene expression in infected cells; 1. global inhibition of protein translation, 2. degradation of cytosolic cellular transcripts and 3. specific translation inhibition of newly transcribed mRNAs. Disruption of cellular protein production using these three components may represent a multi-pronged mechanism that synergistically acts to suppress the host antiviral response (Figure 5C). These mechanisms may explain the molecular basis of the potent suppression of IFN response observed in animal models and in severe COVID-19 patients ^15,18^.

Despite differences in protein size and mode of action, NSP1 proteins from alpha- and beta-CoVs suppress host gene expression ^21,22,46^, making Nsp1 a central host shutoff factor of CoVs. Indeed, several studies focused on SARS-CoV-2 encoded NSP1, showing it binds to the 40S ribosomal subunit, leading to reduction in mRNA translation both in vitro and in cells ^24–28^. Still, how the virus can overcome the NSP1-mediated block of translation and how NSP1 effects come into play during infection remained unclear. We reveal here that in resemblance to what had been described for SARS-CoV NSP1, SARS-CoV-2 NSP1 induces shutdown of host protein translation by two mechanisms: first, it stalls canonical mRNA translation as was reported by others ^24–28,41^ leading to general reduction in the translation capacity of infected cells. Second, NSP1 leads to accelerated cellular mRNA degradation. SARS-CoV NSP1 induces endonucleolytic cleavage and subsequent degradation of host mRNAs and this activity depends on its binding to the 40S ribosome subunit ^21,47^. Our results indicate a similar mechanism operates in SARS-CoV-2 infected cells as we show cytosolic RNAs, specifically ones that are being translated, are more susceptible to SARS-CoV-2-mediated degradation. several studies have shown that mRNAs with viral 5′ UTR are translated more efficiently compared to control 5’UTR in the presence of NSP1 ^24,27,43 26^ but it was further demonstrated that NSP1 inhibits translation of both cellular and a viral 5′ UTR-containing reporter mRNA ^27 26^ implying viral mRNAs do not simply evade translation inhibition in the context of the 5′ UTR sequence. Our measurements from infected cells support these findings as we show viral RNA are not preferentially translated in infected cells. Instead, using a reporter, we show that the viral 5’UTR renders mRNAs refractory to RNA degradation mediated by NSP1. Together these results support a model in which NSP1 acts as a strong inhibitor of translation, tightly binding ribosomes and reducing the pool of available ribosomes that can engage in translation. At the same time ribosome bound NSP1 leads to accelerated degradation of cellular but not of viral mRNAs providing means for viral mRNA to quickly take over and dominate the mRNA pool. This accumulation of SARS-CoV-2 mRNAs explains how infected cells switch their translation towards viral mRNAs. In support of this model, we show that within 8 hours 88% of the mRNAs in infected cells is viral. Notably, interactions between SARS-CoV NSP1 and the viral 5′ UTR prevent NSP1-induced RNA cleavage ^47^, indicating that likely similar mechanisms are applied by SARS-CoV and SARS-CoV-2 viruses.

On top of the general translation reduction and cellular mRNA degradation which are likely mediated by NSP1, we also observed that in mRNAs that are being induced in response to infection there is no corresponding increase in footprints, indicating these newly generated transcripts are less likely to engage with ribosomes. Since mRNAs that are induced in infection are enriched in cytokines and IFN induced genes, this distinctive mechanism may provide additional explanation why infected cells fail to mount an efficient anti-viral response. Although we do not know the underlying molecular mechanism, one appealing hypothesis is viral inhibition of cellular mRNA nuclear export. Since SARS-CoV-2 replicates in the cytoplasm, inhibiting the nuclear export of mRNAs can provide unique advantages as it will specifically inhibit cellular mRNA translation and more explicitly it will lead to suppression of the host’s antiviral response genes which are transcriptionally induced and therefore fully depend on efficient nuclear export. ORF6 was recently shown to co-purify with host mRNA export factors ^48^ and by over expression it disrupts nucleocytoplasmic export ^29^, providing a candidate that may explain the phenotype we observe in infection. Another possibility is that NSP16, which was recently shown to inhibit cellular RNA splicing ^24^, is driving this phenotype as interference with splicing will prevent efficient nuclear export. Indeed, we show here that infection leads to increased levels of intronic reads in many cellular transcripts. Although our analysis reveals some of this signature is attributed to accelerated degradation of mature cellular transcripts it is likely there is also a processing and splicing defect that leads to more intronic reads and eventually aberrant export.

Overall, our study provides an in-depth picture of how SARS-CoV-2 efficiently interferes with cellular gene expression, leading to shutdown of host protein production using a multipronged strategy.

## Acknowledgements

We thank Stern-Ginossar lab members for providing valuable feedback. We thank Nevan Krogan for the SARS-CoV-2 ORFs expression plasmids, Ghil Jona and Weizmann Bacteriology and Genomic Repository Units for technical assistance, Shay Weiss for support and guidance in assays involving SARS-CoV-2. This study was supported by Miel de Botton. Work in the Stern-Ginossar lab is supported by a European Research Council starting grant (StG-2014-638142). N.S-G is an incumbent of the Skirball Career Development Chair in New Scientists and is a member of the European Molecular Biology Organization (EMBO) Young Investigator Program. The authors declare no competing interests.

## Methods

### Cells and viruses

Calu3 cells (ATCC HTB-55) were cultured in 6-well or 10cm plates with RPMI supplemented with 10% fetal bovine serum (FBS), MEM non-essential amino acids, 2mM L-Glutamine, 100Units/ml Penicillin and 1% Na-pyruvate. Monolayers were washed once with RPMI without FBS and infected with SARS-CoV-2 virus, at a multiplicity of infection (MOI) of 3, in the presence of 20 μg per ml TPCK trypsin (Thermo scientific). Plates were incubated for 1 hour at 37°C to allow viral adsorption. Then RPMI medium supplemented with 2% fetal bovine serum, MEM non-essential amino acids, L glutamine and penicillin-streptomycin-Nystatin at 37°C, 5% CO2 was added to each well. SARS-CoV-2 BavPat1/2020 Ref-SKU: 026V-03883 was kindly provided by C. Drosten, Charité–Universitätsmedizin Berlin, Germany. It was propagated (5 passages) and tittered on Vero E6 cells and then sequenced ^9^ before it was used. Handling and working with SARS-CoV-2 virus was conducted in a BSL3 facility in accordance with the biosafety guidelines of the Israel Institute for Biological Research. The Institutional Biosafety Committee of Weizmann Institute approved the protocol used in these studies.

### Preparation of ribosome profiling and RNA sequencing samples

For RNA-seq, cells were washed with PBS and then harvested with Tri-Reagent (Sigma-Aldrich), total RNA was extracted, and poly-A selection was performed using Dynabeads mRNA DIRECT Purification Kit (Invitrogen) mRNA sample was subjected to DNAseI treatment and 3’ dephosphorylation using FastAP Thermosensitive Alkaline Phosphatase (Thermo Scientific) and T4 PNK (NEB) followed by 3’ adaptor ligation using T4 ligase (NEB). The ligated products used for reverse transcription with SSIII (Invitrogen) for first strand cDNA synthesis. The cDNA products were 3’ ligated with a second adaptor using T4 ligase and amplified for 8 cycles in a PCR for final library products of 200-300bp. For Ribo-seq libraries, cells were treated with 100μg/mL CHX for 1 minute. Cells were then placed on ice, washed twice with PBS containing 100μg/mL CHX, scraped from 10cm plates, pelleted and lysed with lysis buffer (1% triton in 20mM Tris 7.5, 150mM NaCl, 5mM MgCl_2_, 1mM dithiothreitol supplemented with 10 U/ml Turbo DNase and 100μg/ml cycloheximide). After lysis samples stood on ice for 2h and subsequent Ribo-seq library generation was performed as previously described ^50^. Briefly, cell lysate was treated with RNAseI for 45min at room temperature followed by SUPERase-In quenching. Sample was loaded on sucrose solution (34% sucrose, 20mM Tris 7.5, 150mM NaCl, 5mM MgCl2, 1mM dithiothreitol and 100μg/ml cycloheximide) and spun for 1h at 100K RPM using TLA-110 rotor (Beckman) at 4c. Pellet was harvested using TRI reagent and the RNA was collected using chloroform phase separation. For size selection, 15uG of total RNA was loaded into 15% TBE-UREA gel for 65min, and 28-34 footprints were excised using 28 and 34 flanking RNA oligos, followed by RNA extraction and ribo-seq protocol^50^

### Sequence alignment, metagene analysis

Sequencing reads were aligned as previously described ^51^. Briefly, linker (CTGTAGGCACCATCAAT) and poly-A sequences were removed and the remaining reads were aligned to the Hg19 and to the SARS-Cov-2 genome (Genebank NC_045512.2) with 3 changes to match the used strain (BetaCoV/Germany/BavPat1/2020 EPI_ISL_406862), 241:C−>T, 3037:C−>T, 23403:A−>G]. Alignment was performed using Bowtie v1.1.2 ^52^ with maximum two mismatches per read. Reads that were not aligned to the genome were aligned to the transcriptome and to SARS-CoV-2 junctions that were recently annotated ^44^. The aligned position on the genome was determined as the 5’ position of RNA-seq reads, and for Ribo-seq reads the p-site of the ribosome was calculated according to read length using the off-set from the 5’ end of the reads that was calculated from canonical cellular ORFs. The offsets used are +12 for reads that were 28-29 bp and +13 for reads that were 30-33 bp. Reads that were in different length were discarded. In all figures presenting ribosome densities data, all footprint lengths (28-33bp) are presented.

Junctions spanning reads were quantified using STAR 2.5.3a aligner ^53^, with running flags as suggested by ^44^, to overcome filtering of non-canonical junctions. Reads aligned to multiple locations were discarded ^44^.

For the metagene analysis only genes with more than 10 reads were used. For each gene, normalization was done to its maximum signal and each position was normalized to the number of genes contributing to the position.

### Filtering of genes, quantification and RPKM normalization

For analysis of cellular genes, the genes were filtered according to the number of reads as follows. The number of reads on a gene in each replicate, from at least one of the conditions (uninfected or 8hr) had to be greater than 50 for the mRNA and greater than 25 for the FP. For heatmaps and clustering, gene filtering was done according to mRNA as described above, and genes with zero reads in any of the samples (mRNA and FP of all time points) were excluded. Histone genes were excluded. Read coverage on genes was normalized to units of RPKM in order to normalize for gene length and for sequencing depth. For analysis comparing host and viral genes, the RPKM was calculated based on the total number of reads of both the host and the virus. For analysis concentrating on cellular genes alone, the RPKM was calculated based on the total number of host reads, and the RPKM values were further scaled to keep the number of total reads equal across samples. The scale factors for RNA-seq were calculated from the ratio of the host mRNA reads to total reads, including viral, rRNA and tRNA reads. in the total RNA sequencing (without polyA selection), and the scale factors for footprints were calculated from the fraction of host reads from the total aligned ribosome profiling reads, including viral and host reads.

Since the viral RNAs are widely overlapping, RNA-seq RPKM levels of viral genes were computed with deconvolution as detailed here. First, values for each gene were calculated by subtracting the RPKM of an ORF from the RPKM of the ORF located just upstream of it in the genome. Then, for subgenomic RNAs, leader-body junction counts were counted based on STAR alignment number of uniquely mapped reads crossing the junction. Finally, based on the correlation between the deconvoluted RPKM and junction abundance of the subgenomic RNAs, the RPKM levels of all viral RNAs was estimated. To quantify the translation levels of non-canonical viral ORFs ORF-RATER was used ^49^.

### Quantification of intronic reads

To avoid biases from intron read count, genes without introns, or where at least one of the introns is overlapping with an exon of another gene were excluded. In addition, genes with low number of reads ( < 20 on the exons, < 2 on the introns) were ignored. The number of reads on exons and introns was normalized by the total length of the exons and introns respectively to get quantification proportional to the number of molecules. Finally, the normalized number of reads on the introns was calculated as percentage of the normalized number of reads on the exons. Statistical significance (in figure 3E) was tested using a paired t-test on the log values of the percentage (with offset of 0.1 to overcome zero values).

### Comparison of cytosolic lncRNAs and mitochondrial mRNAs

The list of cytosolic lncRNAs is based on ^41^. The list of 14 lncRNAs (with number of reads >= 50) that were used in the analysis are given in supplementary table 2. The lncRNAs were compared to protein coding genes, and mitochondrial encoded mRNAs were compared to nuclear encoded mRNAs. Statistical significance was tested using Wilcoxon rank test on the log fold change (compared to uninfected) of the RPKM values.

### Protein synthesis measurement using O-Propargyl Puromycin (OPP)

OPP assay (OPP, Thermo Fisher Scientific) was carried out following the manufacturer’s instructions. Briefly, cells were collected following treatment with 10μM O-Propargyl Puromycin for 30 minutes at 37 ֯C. The cells were then fixed for 15 minutes in 3.7% formaldehyde, and permeabilized in 0.1% Triton X-100 for 15 minutes. OPP was then fluorescently labeled by a 30 minute incubation in Click-iT® Plus OPP reaction cocktail with Alexa Fluor®594 picolyl azide (Thermo Fisher Scientific). Cells were analyzed using BD LSRII flow cytometer.

### Pathway enrichment analysis

Enrichment analysis of cellular pathways in specific gene clusters was done with PANTHER version 15.0, with default settings and the PANTHER pathways data set ^54,55^.

### Reporter assay

pLVX-EF1alpha-SARS-CoV-2-nsp1-2XStrep-IRES-Puro and pLVX-EF1alpha-SARS-CoV-2-nsp2-2XStrep-IRES-Puro were kindly provided by Nevan Krogan, University of California, San Francisco.

The viral 5’UTR was constructed based on nucleotides 4-265 of the reported sequence of SARS-CoV-2 isolate Wuhan-Hu-1 (NC_045512.2) by sequential annealing of DNA oligonucleotides (IDT, 5’UTR oligo_1-5 listed in the table below). The coding sequence for the first 12 amino acids (aa) of ORF1a as well as the AcGFP homology region were added to the 5’UTR by two PCR amplifications. For the 5’UTR-less control plasmid, the 12 aa region with AcGFP homology was ordered from Sigma-Aldrich as oligonucleotides. The viral 5’UTR with the 12 aa region and the 12 aa region on its own were cloned into pAcGFP1-C1 (Takara Biotech) using restriction-free cloning. The entire expression cassette from the promoter to the poly-A site was amplified and cloned into pDecko-mCherry (Addgene plasmid #78534) using restriction-free cloning. Primers for PCR amplification of fragments were ordered from Sigma-Aldrich. All primers and oligonucleotides used for cloning are listed in the table below.

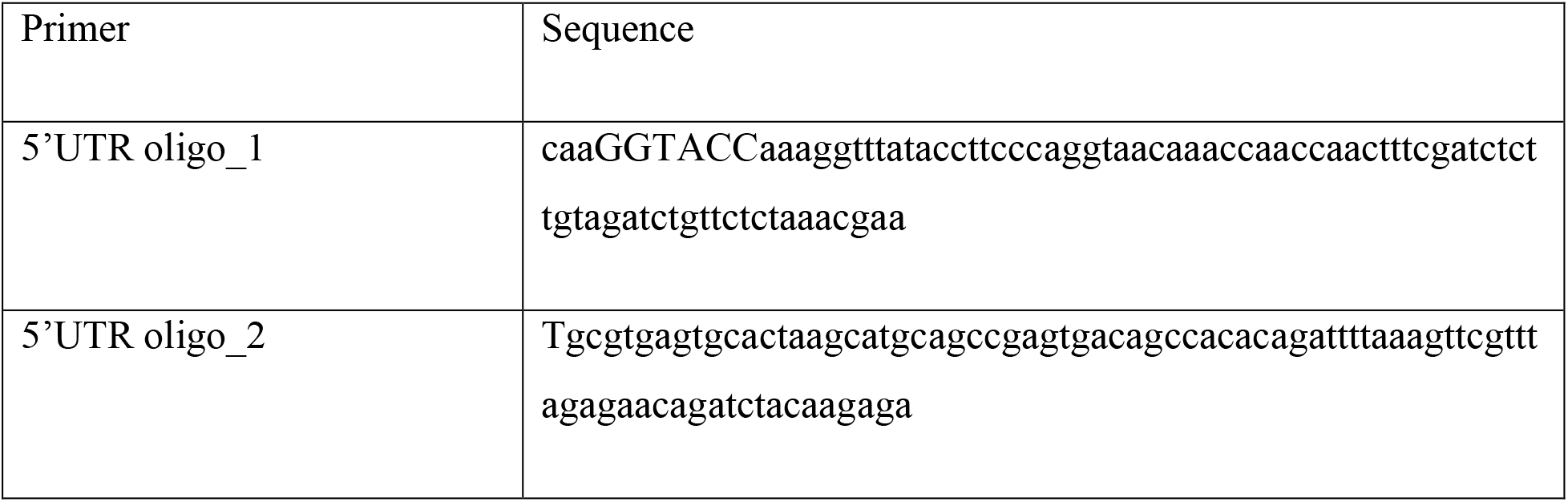
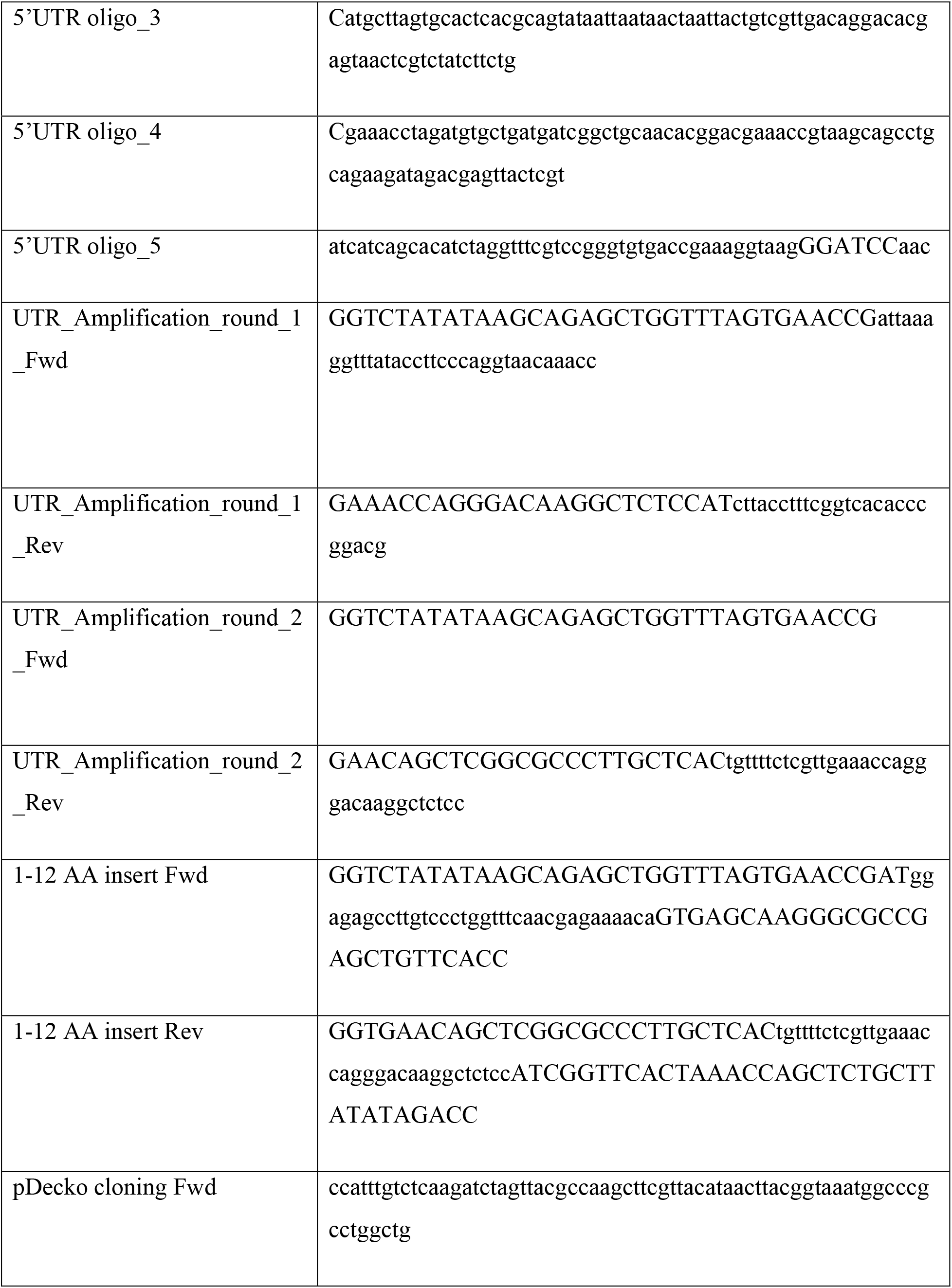
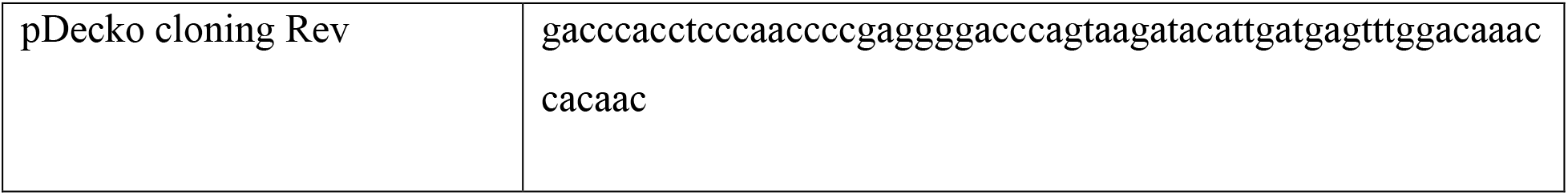

293T cells were transfected using JetPEI (Polyplus-transfection) following the manufacturer’s instructions. Cells were assayed 24 hours post transfection by flow cytometry on a BD Accuri C6 flow cytometer and imaged on a Zeiss AxioObserver Z1 wide-field microscope using a X20 objective and Axiocam 506 mono camera.

### Quantitative real-time PCR analysis

Total RNA was extracted using Tri-Reagent (Sigma) and polyA selected using the Dynabeads mRNA DIRECT purification kit (Thermo Fisher Scientific) following the manufacturer’s instructions. cDNA was prepared using qScript cDNA Synthesis Kit (Quanta Biosciences) following the manufacturer’s instructions. Real time PCR was performed using the SYBR Green PCR master-mix (ABI) on the QuantStudio 12K Flex (ABI) with the following primers (forward, reverse):

ActinB (GTCATTCCAAATATGAGATGCGT, GCTATCACCTCCCCTGTGTG)

AcGFP (TGACCCTGAAGTTCATCTGC, GAAGTCGTGCTGCTTCATGT)

mCherry (ACCGCCAAGCTGAAGGTGAC, GACCTCAGCGTCGTAGTGGC)

GFP and mCherry mRNA levels were calculated relative to the human ActinB transcript.

