## Supplementary figures for "SARS-CoV-2 utilizes a multipronged strategy to suppress host protein synthesis"

Supplementary Figure 1

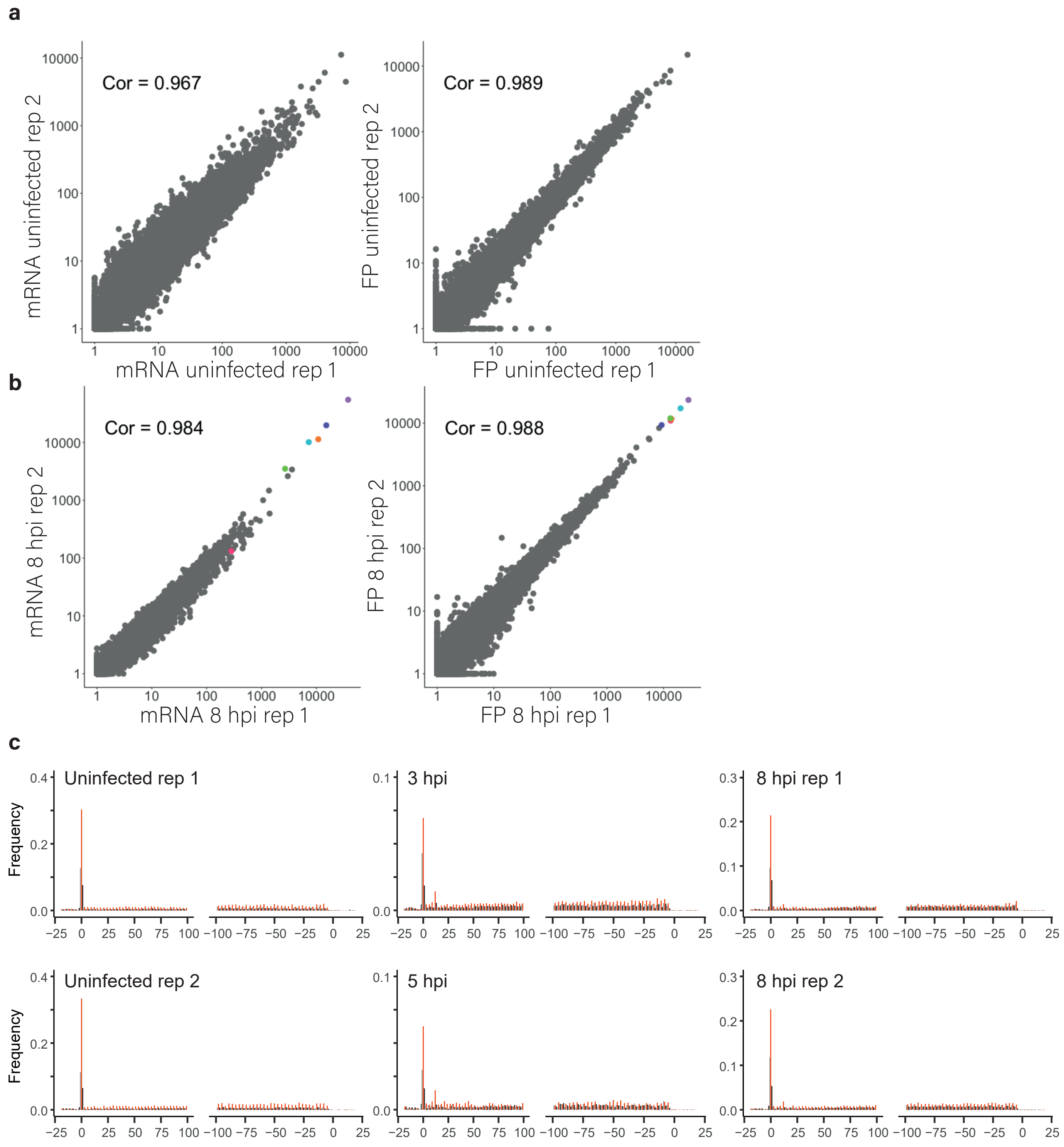


**Figure S1:** data replicability and quality
(**A and B**) Scatter plots depicting gene densities derived from our two independent biological replicates for mRNA **(A)** and footprints **(B)**, demonstrating reproducibility between our replicates. Pearson's R values are presented. **(C)** Metagene analysis of read densities around the start codon of protein coding genes as measured by our ribosome profiling libraries at different time points post infection. The X axis shows the nucleotide position relative to the start codon. The ribosome densities are shown with different colors indicating the three relative frames. The translated frame labeled in red (red, frame 0; black, frame +1; grey, frame +2). All libraries show 3bp periodicity.

Figure S2

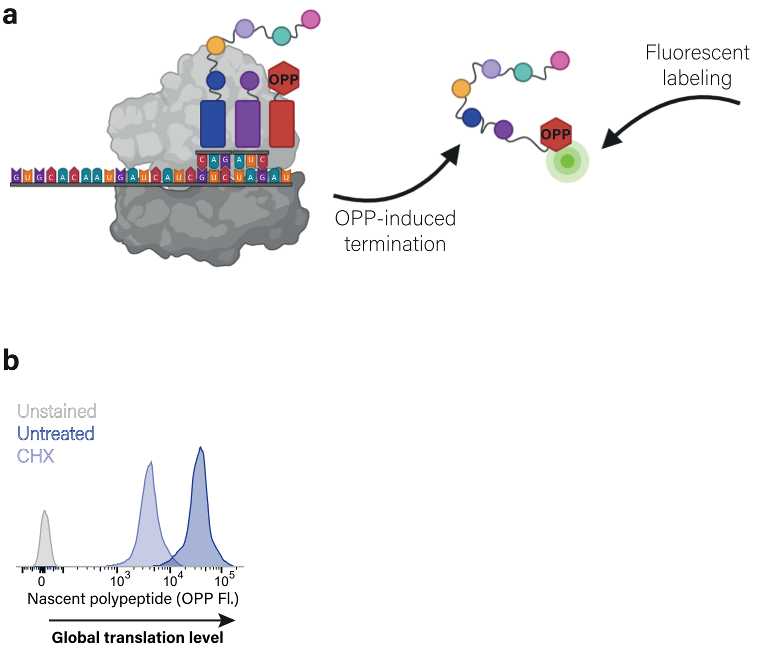


**Figure S2**: measurement of protein synthesis using the OPP assay
**(A)** Schematic depiction of labeling and detection of nascent protein synthesis by O-propargyl-puromycin (OPP) incorporation followed by florescent labeling. OPP is efficiently incorporated into newly translating proteins, releases the polypeptides and terminates translation. Following fixation, OPP is fluorescently labeled using Click reaction and can then be measured by flow cytometry. **(B)** Protein synthesis measurement by flow cytometry of untreated 293T cells and 293T cells treated with cycloheximide leading to efficient translation inhibition. Unlabeled cells are shown as control.

Supplementary figure 3

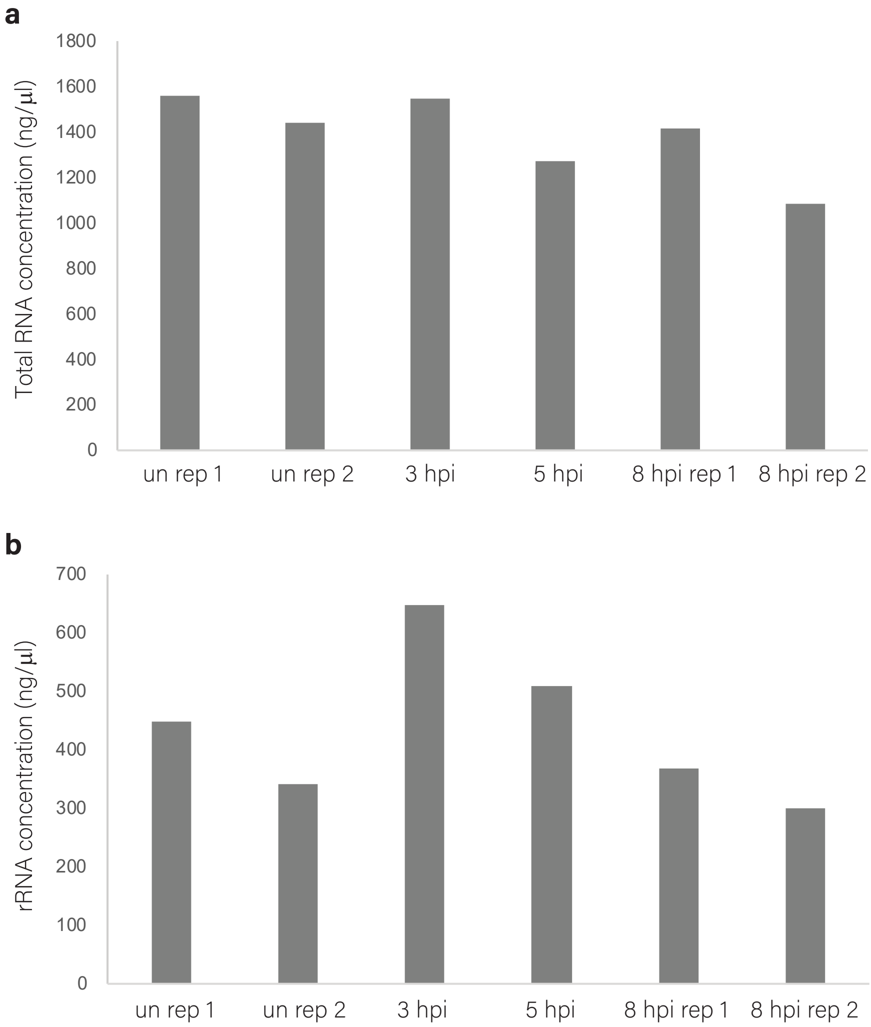


**Figure S3**: total RNA and ribosomal RNA measurements from infected and uninfected Calu3 cells.
(**A**) Total RNA was measured in uninfected Calu3 cells and in cells at 3, 5 and 8 hpi by Qubit Fluorometer. **(B)** rRNA concentration was measured in uninfected Calu3 cells and in cells at 3, 5 and 8 hpi by measuring the concentration in the 18S and 28S ribosomal RNA peaks using a TapeStation system.

Supplementary figure 4

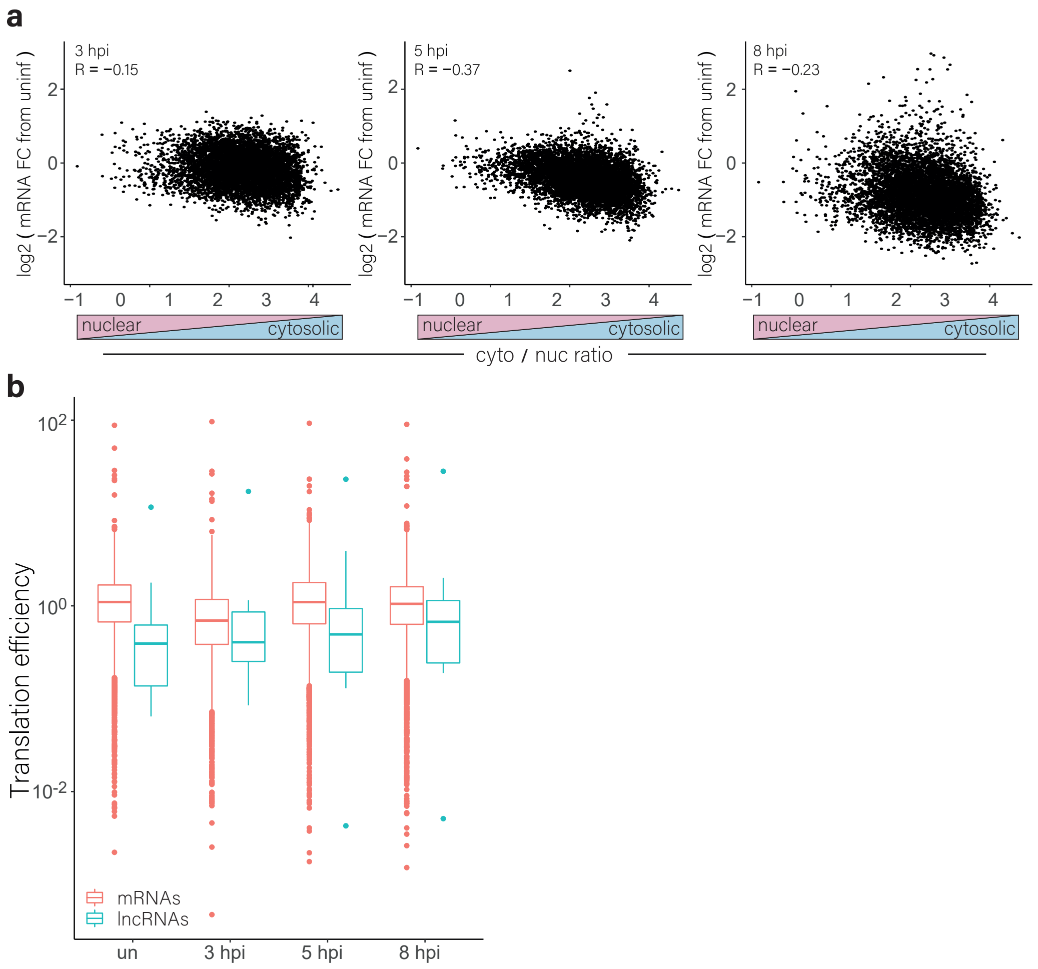


**Figure S4**: cytosolic lncRNAs expression levels
**(A)** Scatter plots depicting nuclear to cytosol ratio of cellular transcripts as was calculated from [41] relative to changes in transcript levels between uninfected cells and cells at the different time points during SARS-CoV-2 infection. **(B)** The translation efficiency of cytosolic lncRNAs and protein coding mRNAs in uninfected cells and at different time points after infection.

Supplementary figure 5

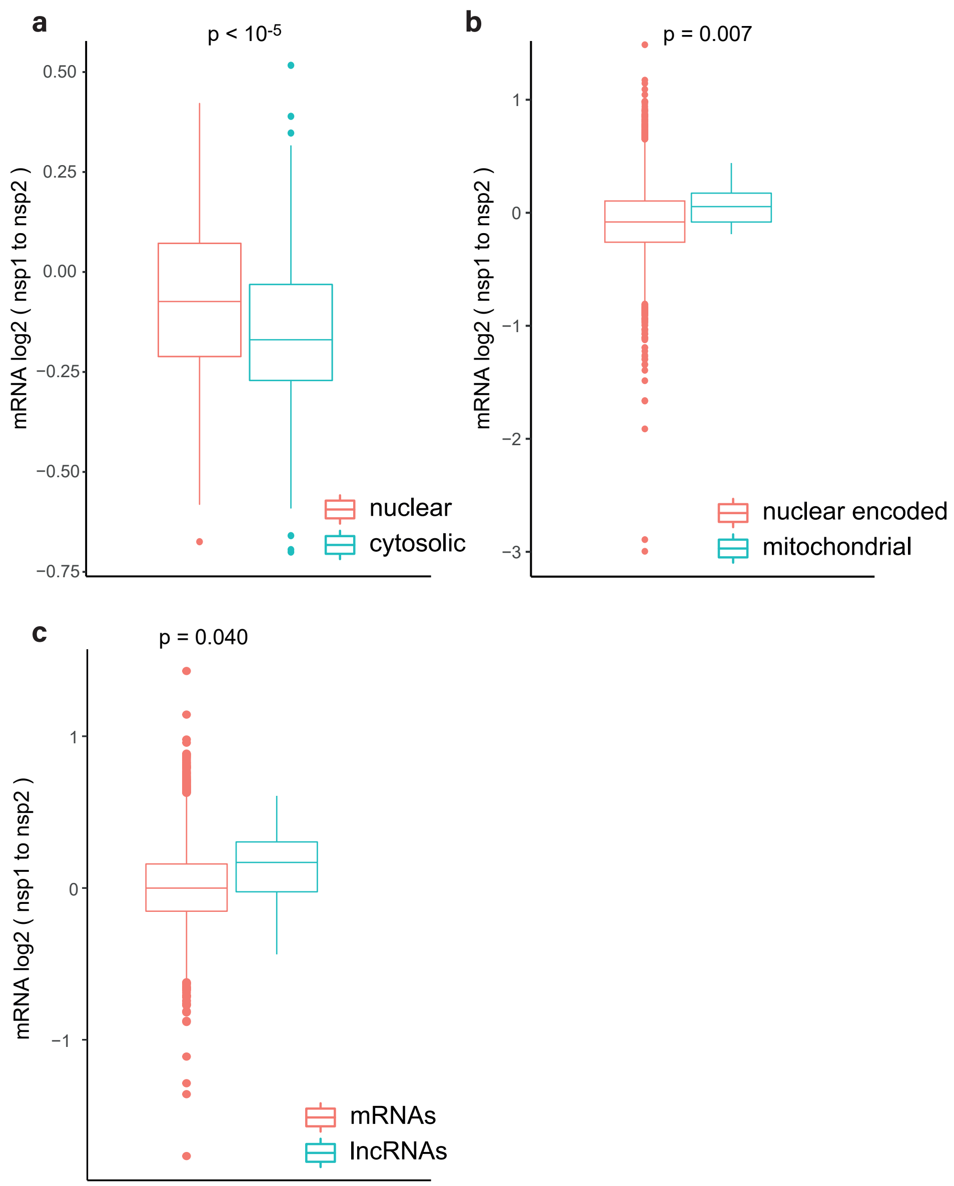


**Figure S5:** NSP1 effect on RNA levels
**(A)** RNAs were grouped to two bins based on their cytosol to nucleus localization ratio [41]. Presented in the fold change in RNA expression in cells transfected with NSP1 relative to cells transfected with NSP2. **(B)** The change in RNA expression of nuclear encoded or mitochondrial encoded RNAs in cells transfected with NSP1 relative to cells transfected with NSP2. **(C)** The change in RNA expression of cytosolic lncRNAs and protein coding mRNAs in cells transfected with NSP1 relative to cells transfected with NSP2. RNA-seq data of NSP1 and NSP2 transfected cells is from [42].

Supplementary figure 6

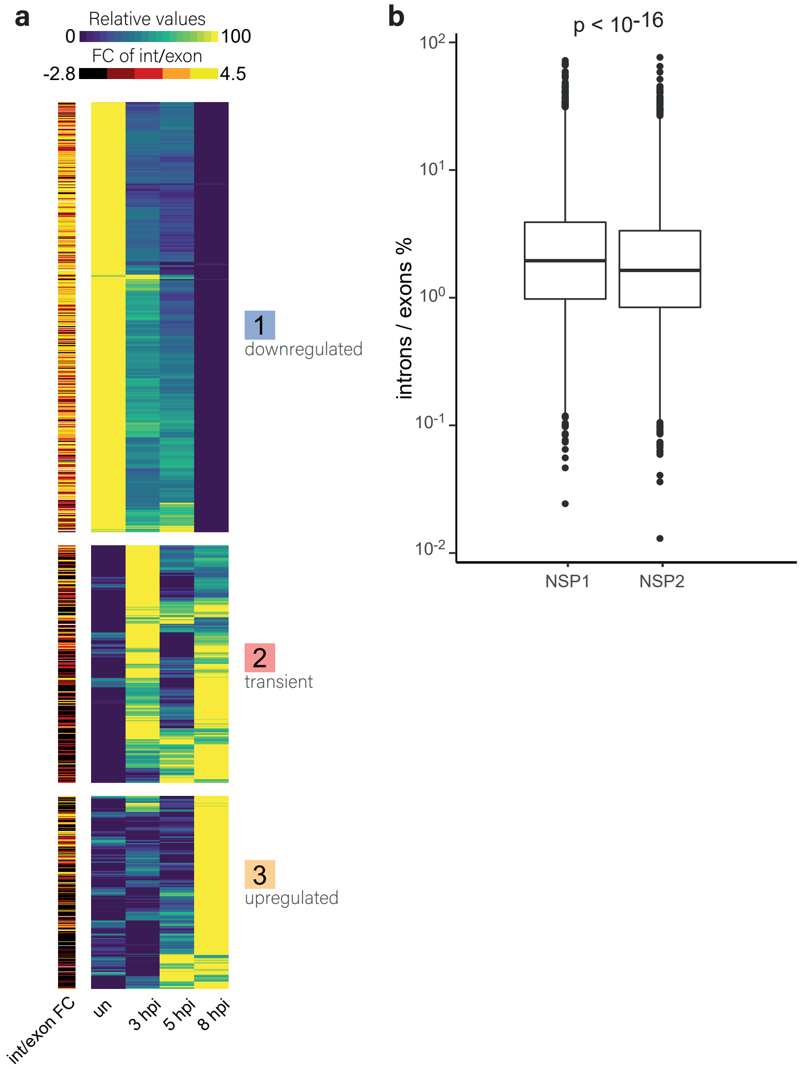


**Figure S6:** increase in intronic reads
**(A)** Heat map presenting relative mRNA and footprints expression of well-expressed human transcripts that showed the most significant changes in their mRNA levels at 8 hpi relative to uninfected, across time points during SARS-CoV-2 infection (as presented in Figure 1G). Three main clusters are marked on the right. Presented on the right is the fold change in the ratio of intron to exon reads for each transcript in uninfected compared to 8hpi. **(B)** Box plots presenting the ratio of intronic to exonic reads for each gene in cells transfected with NSP1 relative to cells transfected with NSP2. RNA-seq data of NSP1 and NSP2 transfected cells is from [42].

Supplementary figure 7

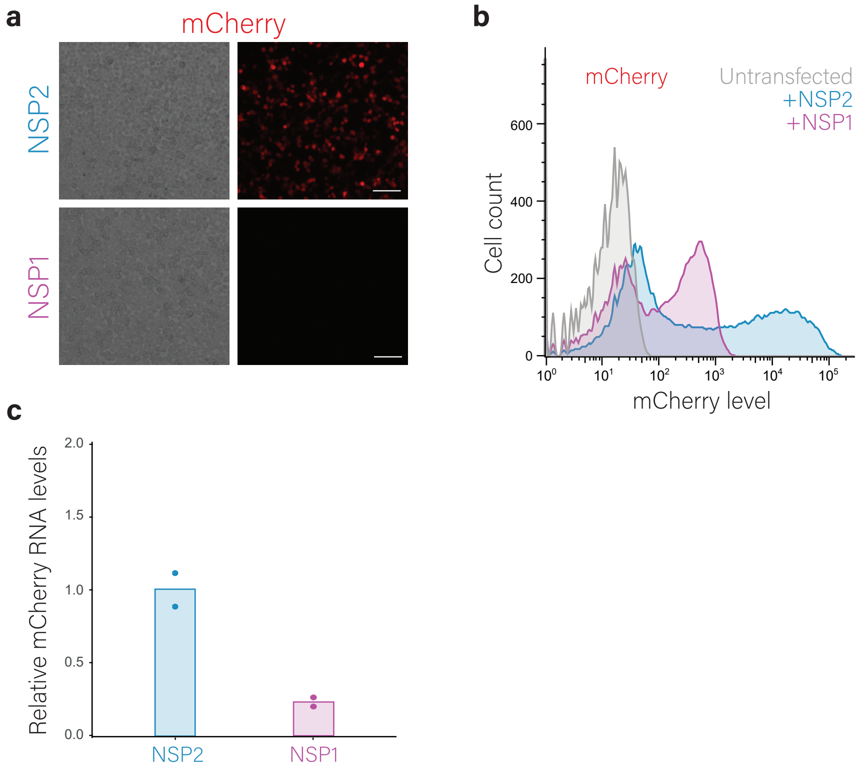


**Figure S7:** NSP1 reporter assay
293T cells were transfected with expression vectors of either NSP1 or NSP2 (as a control) and with the GFP (+) 5'UTR reporter containing an independent mCherry reporter. **(A)** Microscopy images showing mCherry expression of cells co-transfected with NSP2 (top) or NSP1 (bottom) together with GFP (+) 5'UTR/mCherry reporter. **(B)** Flow cytometry analysis of the mCherry expression from cells co-transfected with NSP1 or NSP2 together with GFP (+) 5'UTR/mCherry reporter. **(C)** Relative mCherry RNA levels in cells expressing NSP1 or NSP2 together with GFP (+) 5'UTR/mCherry reporter as measured by quantitative RT-PCR. Data points show measurement of biological replicates. Shown is one representative experiment out of two performed.

Supplementary figure 8

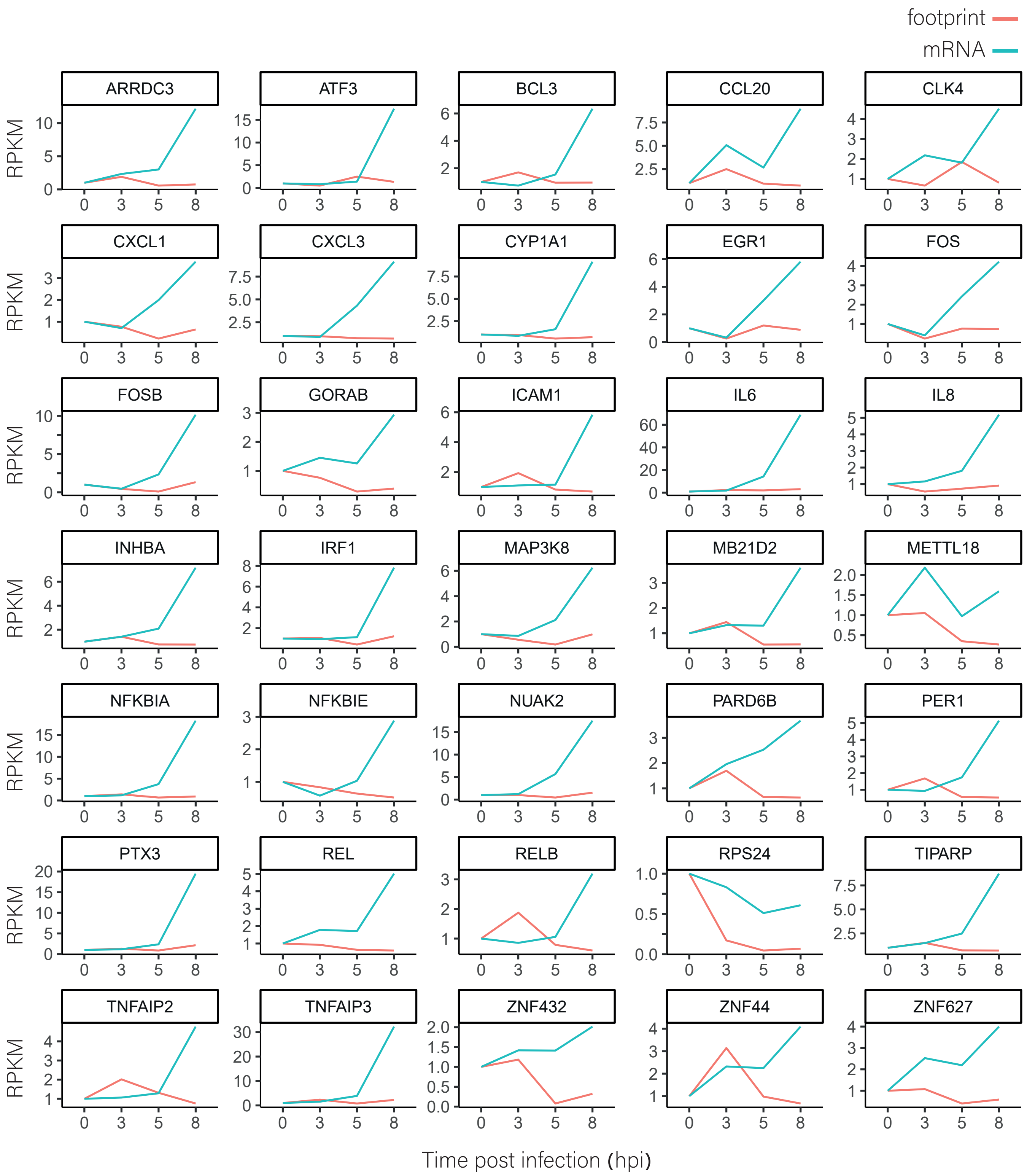


**Figure S8:** expression kinetic of TE reduced genes
mRNA and footprint levels of the indicated genes in uninfected cells and at 3, 5 and 8 hpi. Presented are genes that were significantly reduced in their relative TE.
